## Supplementary information for "A causal role of the NMDA receptor in recurrent processing during perceptual inference: NMDA receptor antagonist memantine selectively improves EEG decoding of the Kanizsa illusion"

This research was supported by a grant from the H2020 European Research Council (ERC STG 715605, SVG).

Participant exclusion criteria were:

- Allergy for memantine or one of the inactive ingredients of these products;
- (History of) psychiatric treatment;
- First-degree relative with (history of) schizophrenia or major depression;
- (History of) clinically significant hepatic, cardiac, obstructive respiratory, renal, cerebrovascular, metabolic or pulmonary disease, including, but not limited to fibrotic disorders;
- Claustrophobia;
- Regular usage of medicines (antihistamines or occasional use of paracetamol);
- (History of) neurological disease;
- (History of) epilepsy;
- Abnormal hearing or (uncorrected) vision;
- Average use of more than 15 alcoholic beverages weekly;
- Smoking
- History of drug (opiate, LSD, (meth)amphetamine, cocaine, solvents, cannabis, or barbiturate) or alcohol dependence;
- Any known other serious health problem or mental/physical stress;
- Used psychotropic medication, or recreational drugs over a period of 72 hours prior to each test session,
- Used alcohol within the last 24 hours prior to each test session;
- (History of) pheochromocytoma.
- Narrow-angle glaucoma;
- (History of) ulcer disease;
- Galactose intolerance, Lapp lactase deficiency or glucosegalactose malabsorption.

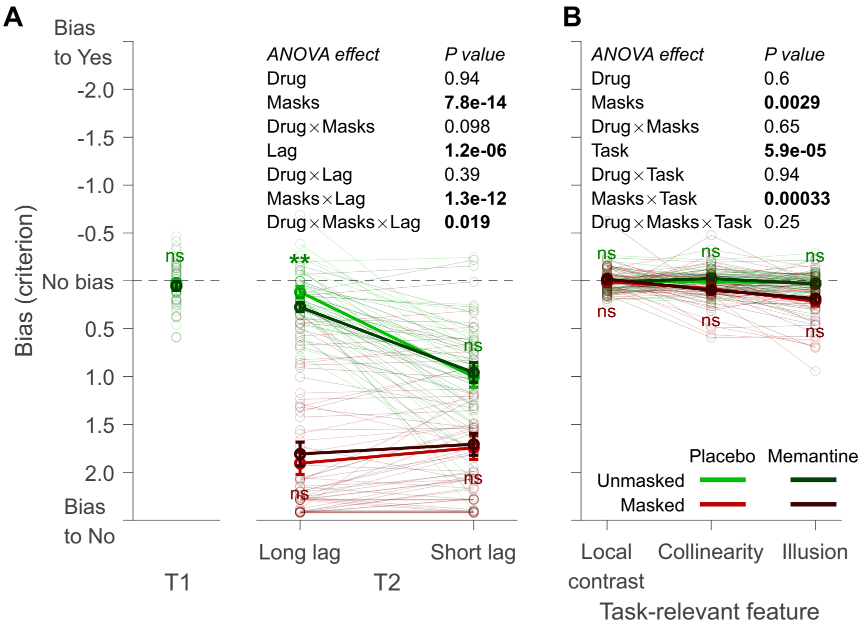


**Figure S1. Bias.** The participants' propensity to indicate the presence of (A) the Kanizsa illusion in experiment 1 and (B) the task-relevant feature in experiment 2. Ns is not significant (*P*>0.05). Error bars are mean ± standard error of the mean. Individual data points are plotted using low contrast. ***P*=0.007.

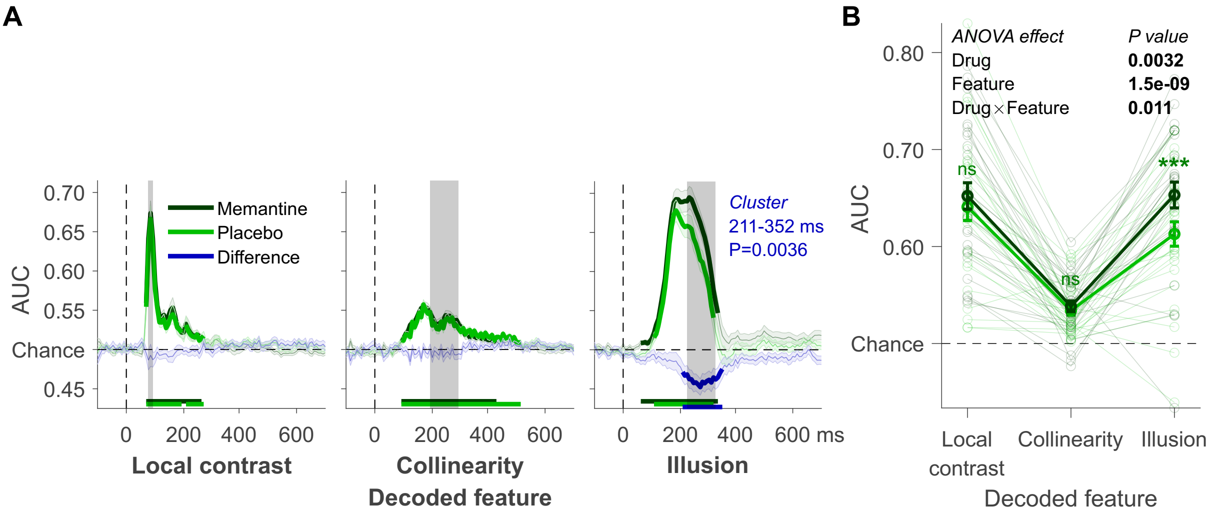


**Figure S2. Diagonal decoding applied to the T1 data.** (A) Local contrast, collinearity, and Kanizsa illusion decoding. These three visual features' time windows: 78-94, 192-292, and 223-323 ms, respectively. Mean decoding performance, area under the receiver operating characteristic curve (AUC), over time ± standard error of the mean (SEM). Thick lines differ from chance: P<0.05, cluster-based permutation test. (B) Mean AUC for every time window. Error bars are mean ± SEM. Individual data points are plotted using low contrast. Ns is not significant (*P*>0.05). ****P*<0.001.

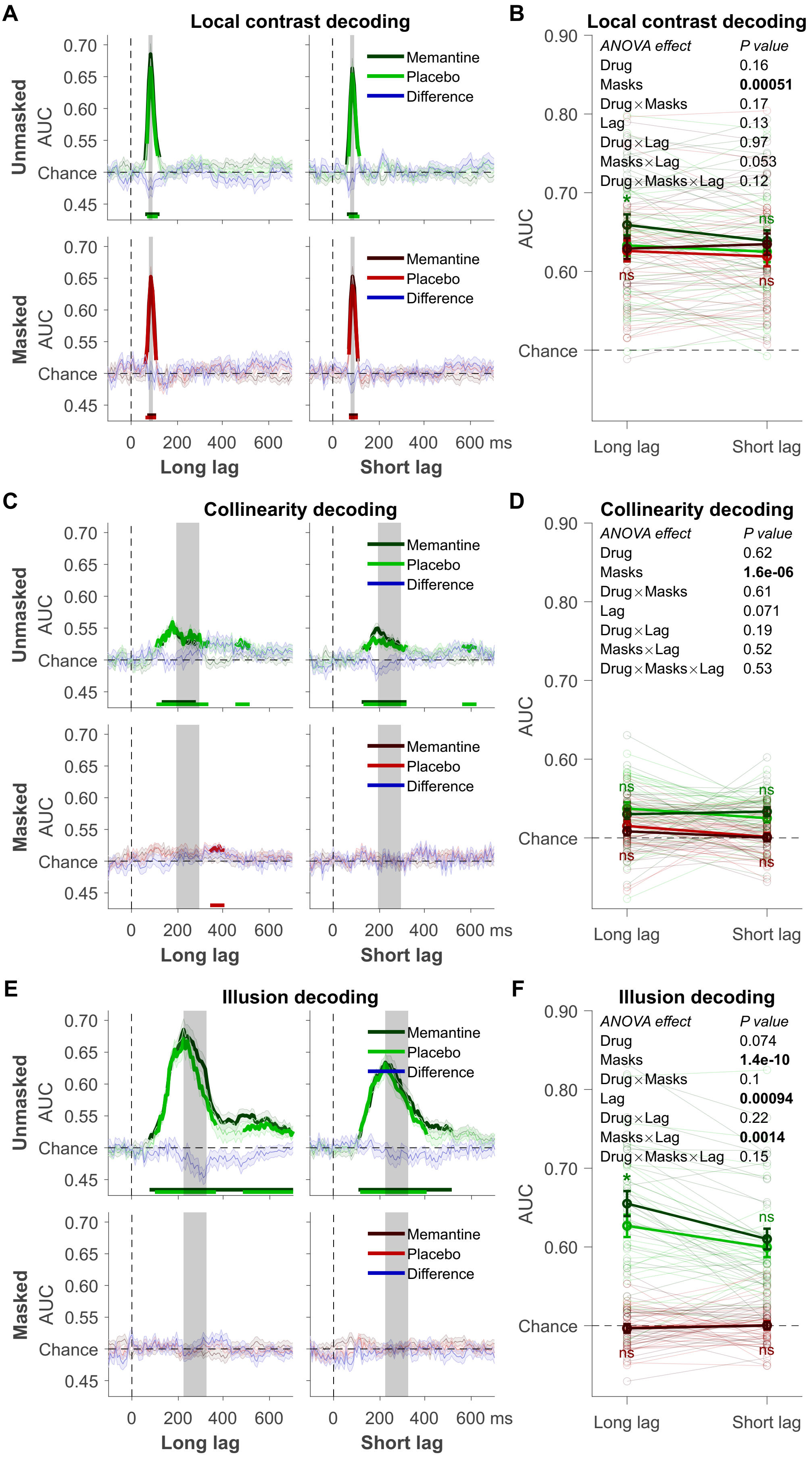
A
Local contrast decoding
Unmasked
Masked
AUC
0.70
0.65
0.60
0.55
Chance
0.45
Memantine
Placebo
Difference
0 200 400 600
0 200 400 600 ms
Long lag
Short lag
B
Local contrast decoding
ANOVA effect	P value
Drug 0.16
Masks 0.00051
Drug×Masks 0.17
Lag 0.13
Drug×Lag 0.97
Masks×Lag 0.053
Drug×Masks×Lag 0.12
0.90
0.80
0.70
0.60
Chance
Long lag
Short lag
C
Collinearity decoding
D
Collinearity decoding
ANOVA effect	P value
Drug 0.62
Masks 1.6e-06
Drug×Masks 0.61
Lag 0.071
Drug×Lag 0.19
Masks×Lag 0.52
Drug×Masks×Lag 0.53
E
Illusion decoding
F
Illusion decoding
ANOVA effect	P value
Drug 0.074
Masks 1.4e-10
Drug×Masks 0.1
Lag 0.00094
Drug×Lag 0.22
Masks×Lag 0.0014
Drug×Masks×Lag 0.15
ns
*

58

**Figure S3. Diagonal decoding of the T2 data.** (A) Local contrast decoding: mean decoding performance, area under the receiver operating characteristic curve (AUC), over time ± standard error of the mean (SEM). Thick lines differ from chance: $P<0.05$, cluster-based permutation test. (B) Mean AUC for local contrast (rotation) decoding's time window: 78 -94 ms. Error bars are mean ± SEM. Individual data points are plotted using low contrast. Ns is not significant ($P>0.05$). (C) Collinearity decoding: mean AUC over time. (D) Mean AUC for collinearity decoding's time window: 192-292 ms. (E) Kanizsa illusion decoding: mean AUC over time. (F) Mean AUC for Kanizsa illusion's time window: 223-323 ms. *$P=0.019$.

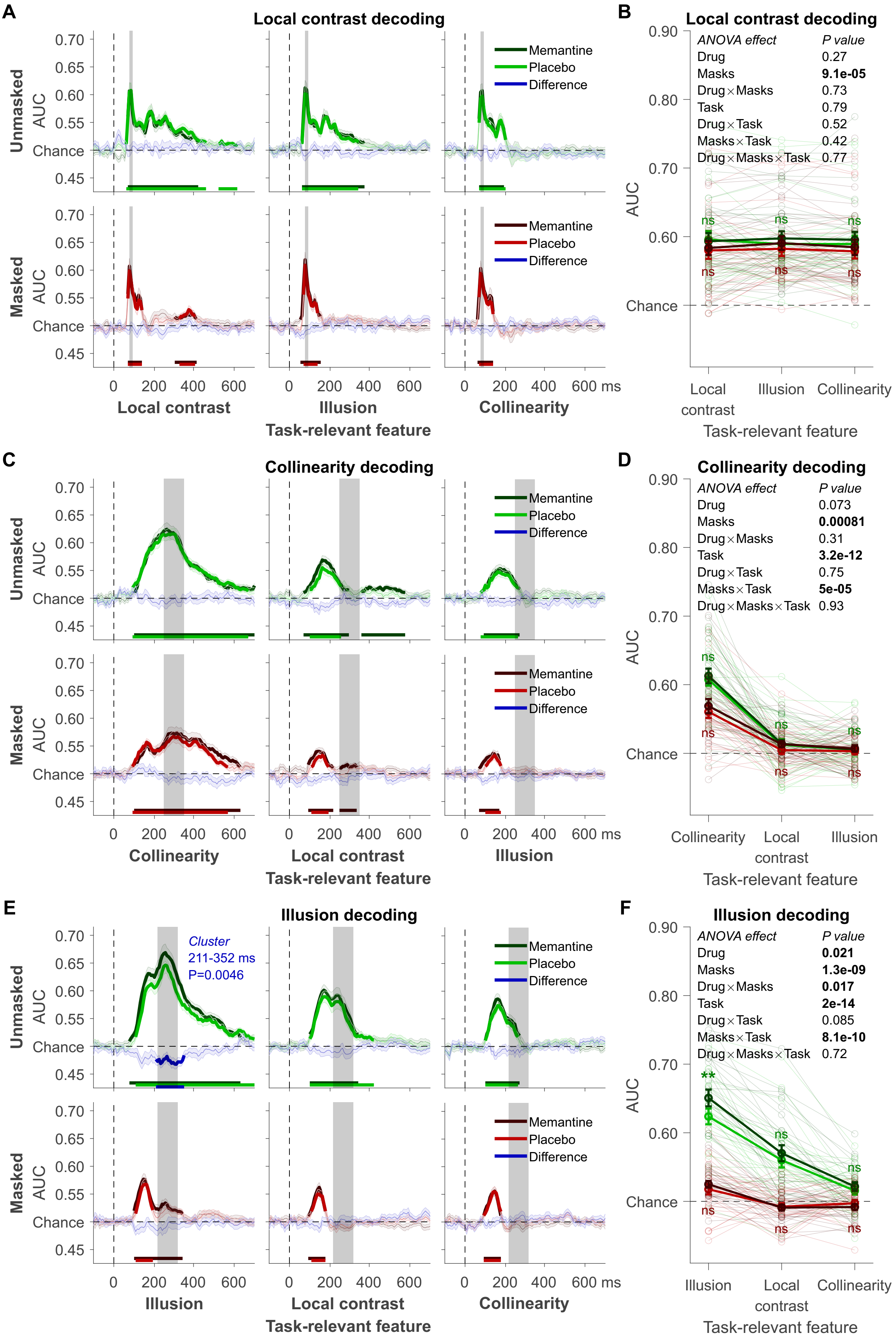
A
Local contrast decoding
Unmasked
Masked
AUC
0.70
0.65
0.60
0.55
Chance
0.45
Memantine
Placebo
Difference
0 200 400 600
0 200 400 600
0 200 400 600 ms
Local contrast
Illusion
Collinearity
Task-relevant feature
B
Local contrast decoding
ANOVA effect P value
Drug 0.27
Masks 9.1e-05
Drug×Masks 0.73
Task 0.79
Drug×Task 0.52
Masks×Task 0.42
Drug×Masks×Task 0.77
0.90
0.80
0.70
0.60
Chance
ns
Local contrast
Illusion
Collinearity
Task-relevant feature
C
Collinearity decoding
Collinearity
Local contrast
Illusion
Task-relevant feature
D
Collinearity decoding
ANOVA effect P value
Drug 0.073
Masks 0.00081
Drug×Masks 0.31
Task 3.2e-12
Drug×Task 0.75
Masks×Task 5e-05
Drug×Masks×Task 0.93
Collinearity
Local contrast
Illusion
Task-relevant feature
E
Illusion decoding
Cluster
211-352 ms
P=0.0046
Illusion
Local contrast
Collinearity
Task-relevant feature
F
Illusion decoding
ANOVA effect P value
Drug 0.021
Masks 1.3e-09
Drug×Masks 0.017
Task 2e-14
Drug×Task 0.085
Masks×Task 8.1e-10
Drug×Masks×Task 0.72
**
Illusion
Local contrast
Collinearity
Task-relevant feature

**Figure S4. Diagonal decoding results of experiment 2.** (A) Local contrast decoding: mean decoding performance, area under the receiver operating characteristic curve (AUC), over time ± standard error of the mean (SEM). Thick lines differ from chance: P<0.05, cluster-based permutation test. (B) Mean AUC for local contrast decoding's time window: 78-94 ms. Error bars are mean ± SEM. Individual data points are plotted using low contrast. Ns is not significant (*P*>0.05). (C) Collinearity decoding: mean AUC over time. (D) Mean AUC for collinearity decoding's time window: 217-317 ms. (E) Kanizsa illusion decoding: mean AUC over time. (F) Mean AUC for Kanizsa illusion's time window: 223-323 ms. ***P*=0.001.

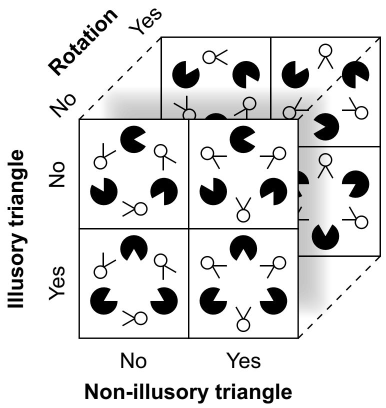


**Figure S5. Stimulus set of experiment 2.**

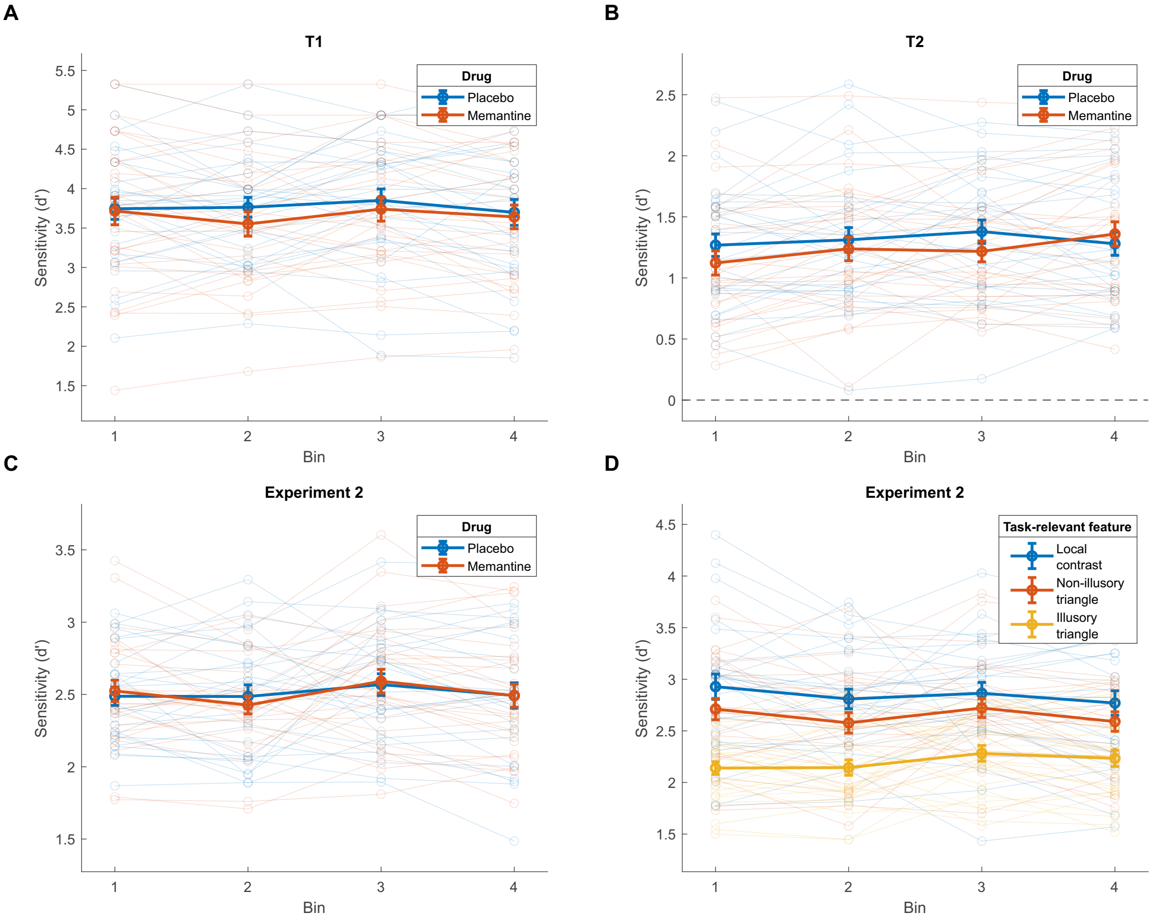


**Figure S6. Behavior over the course of the experiments, divided into four time bins.** (A) Experiment

1: first target (T1). For T1 in the first experiment, there was no interaction between the factors bin and drug (memantine/placebo; $F_{3,84}$=0.89, $P$=0.437). (B) Experiment 1: second target (T2). For T2, we performed a repeated-measures ANOVA with the factors bin, drug, T1-T2 lag (short/long), and masks (present/absent). There was only a trend towards a bin by drug interaction ($F_{3,84}$=2.57, $P$=0.064), reflecting worse performance under memantine in the first three bins and slightly better performance in the fourth bin. The other interactions that include the factors bin and drug factors were not significant (all $P$>0.117). (C) Experiment 2 T2: bin by drug interaction and (D) Experiment 2 T2: bin by task-relevant feature interaction. For the second experiment, we performed a repeated-measures ANOVA with the factors bin, drug, masks, and task-relevant feature (local contrast/collinearity/illusion). None of the interactions that included the bin and drug factors were significant (all $P$>0.219). Taken together, memantine d id not appear to affect Kanizsa illusion detection performance through perceptual learning. Finally, there was no interaction between the factors bin and task-relevant feature ($F_{6,150}$=0.76, $P$=0.547), suggesting that there was no perceptual learning effect specific to Kanizsa illusion detection.
